# Diversity and evolution of the transcriptional regulatory networks of *Pseudomonas* strains revealed using machine learning

**DOI:** 10.64898/2026.07.20.739605

**Authors:** Heera Bajpe, Ying Hefner, Richard Szubin, Jaemin Sung, Bernhard O. Palsson

## Abstract

The genus *Pseudomonas* consists of diverse and ecologically significant species that form close associations with both plants and animals. This genus is widely studied due to the clinically relevant *Pseudomonas aeruginosa*, model plant pathogen *Pseudomonas syringae*, and non-pathogenic, industrially relevant *Pseudomonas putida*. The different metabolic and physiological capabilities of these species are enabled by their unique genetic makeup as well as varying regulatory mechanisms. To study the transcriptional basis for the diversity of the three species, we applied independent component analysis to strain-specific RNA-seq datasets to identify independently modulated gene sets (iModulons) and their condition-specific activity levels. We then mapped iModulons across strains based on their similarity in orthologous gene membership. Through comparison of iModulon gene membership and activities, we find that: (i) iModulons reveal shared and unique regulatory modalities across strains; (ii) unique adaptations in common functions, such as translation and pyoverdine production/uptake, manifest through both differential iModulon gene membership and condition-specific activation states in each strain; (iii) iModulons facilitate comparison of stress responses at the systems level; and (iv) iModulons highlight unique virulence factor enrichment and host-specific adaptations in human and plant pathogens. Altogether, comparing the modularized transcriptomes of the three strains provides unique and comprehensive insights into their differential evolution.

**Importance:** Closely related bacterial species often have vastly different metabolic and physiological capabilities, yet the regulatory mechanisms underlying these adaptations remain poorly understood. Here, we compare the transcriptional regulatory networks of three representative *Pseudomonas* strains through cross-strain iModulon analysis. By comparing both iModulon gene composition and activity across strains, we identify conserved regulatory modules alongside lineage-specific adaptations in functions associated with virulence, translation, iron acquisition, motility, and stress responses. Our results demonstrate that iModulons provide a genome-scale framework for comparing transcriptional regulation across closely related organisms, revealing regulatory innovations that are not apparent from genome comparisons alone. This work establishes a scalable approach for studying the evolution of bacterial transcriptional regulatory networks and the regulatory basis of niche specialization.

## Introduction

*Pseudomonas* is a complex genus containing the largest number of gram-negative bacterial species(1, 2). *Pseudomonas aeruginosa*, *Pseudomonas syringae*, and *Pseudomonas putida* are prominent species of this genus with high genomic similarity, but diverse functionalities(3). *P. aeruginosa* is an opportunistic pathogen that is a leading cause of nosocomial infections in humans(4). *P. syringae* is a phytopathogen responsible for agricultural and economic losses worldwide(3). Finally, *P. putida* is a largely non-pathogenic saprophytic species of industrial relevance(5). While the metabolic and physiological versatility of these pseudomonads is in part due to genomic differences, unique regulatory mechanisms enable survival in diverse niches. These regulatory mechanisms are controlled by the transcriptional regulatory network (TRN). Genome-scale TRN comparison is imperative to furthering our knowledge on the evolution of transcriptional regulation across this lineage.

The TRN is a complex network of regulators and their target genes. TRN characterization is limited by bottom-up approaches that tend to be tedious and expensive(6). However, rapid growth of RNA-seq data and development of scalable data analytic tools has furthered efforts to deconvolute the TRN through top-down approaches. Independent component analysis (ICA), a blind source separation algorithm that identifies statistically independent components in complex datasets, has been proven to successfully untangle bacterial transcriptomes(7–10). When applied to bacterial gene expression profiles, it identifies independently modulated gene sets called iModulons. ICA determines the gene composition of each iModulon and its activity across all conditions in the dataset. iModulons represent groups of genes that are coregulated under certain conditions, thereby corresponding to interpretable regulatory signals. Modularization of bacterial transcriptomes has previously enabled the discovery of regulons, characterization of unknown genes and regulons, and identification of transcriptomic trade-offs(6, 10, 11).

We previously used iModulon analysis to study the TRNs of *P. aeruginosa* PAO1(12, 13), *P. syringae* pv. *tomato* DC3000(14), and *P. putida* KT2440(5). iModulons of *P. aeruginosa* revealed biosynthetic gene cluster (BGC) and multidrug resistance-associated regulation that contribute to virulence. iModulons of *P. syringae* provided insight into microbe-microbe and plant-pathogen interactions. Finally, iModulons of *P. putida* uncovered transcriptional responses relevant to industrial bioprocesses. iModulons from these pseudomonads have also guided metabolic engineering through the transfer of catabolic iModulons across species to confer novel substrate readiness on the recipient strain(15). These studies highlight strain-specific regulatory mechanisms that enable each strain to survive in their unique environments.

iModulons also enable comparison of regulatory mechanisms across strains. A previous study comparing iModulons of seven strains across different genera uncovered conserved and lineage-specific regulatory aspects across the TRNs(16). The application of this methodology to strains belonging to a single genus is expected to reveal common and unique TRN adaptations in closely related strains. Here, we use iModulon analysis to compare the TRNs of *P. aeruginosa* PAO1, *P. syringae* pv. *tomato* DC3000, and *P. putida* KT2440. We first expanded the RNA-seq datasets of each strain by adding samples generated under common conditions. Then, we used ICA to recompute strain-specific iModulon structures.

Using the updated iModulon structures, we mapped iModulons across the strains based on orthologous gene membership and compared TRNs based on structure and activation state. We find that strain-specific adaptations within conserved functions, such as translation and pyoverdine biosynthesis/uptake, include unique iModulon gene membership and activation states. Additionally, motility-associated iModulons reveal strain-specific requirements influenced by varying lifestyles of pathogens and saprophytes. Furthermore, genome-wide antibiotic stress responses vary across strains, with the *P. aeruginosa* TRN having specialized antibiotic stress response components. Our analysis thus details differential adaptation and evolution of TRNs across three *Pseudomonas* strains through the unique lens of iModulons.

## Results

### Reconstruction of iModulon structures

Although a few conditions are roughly comparable across the datasets used in our previous studies on the three strains, there is an overall lack of comparability (Supplementary Note S1). This results in differences in iModulon structures and hinders iModulon activation comparisons under identical conditions across strains. To address this limitation, we expanded each dataset by adding RNA-seq samples generated under common conditions. After quality control, each dataset contained eight newly generated common conditions, including nutrient starvation, metal ion stress, antibiotic stress, and oxidative stress (Supplementary Note S2). The datasets of *P. aeruginosa*, *P. putida*, and *P. syringae* were expanded to 431, 343, and 244 RNA-seq samples, respectively (Supplementary Table S1-3). Principal component analysis reveals that growth conditions largely contribute to the separation between samples (Fig. 1A).

**Figure 1.**
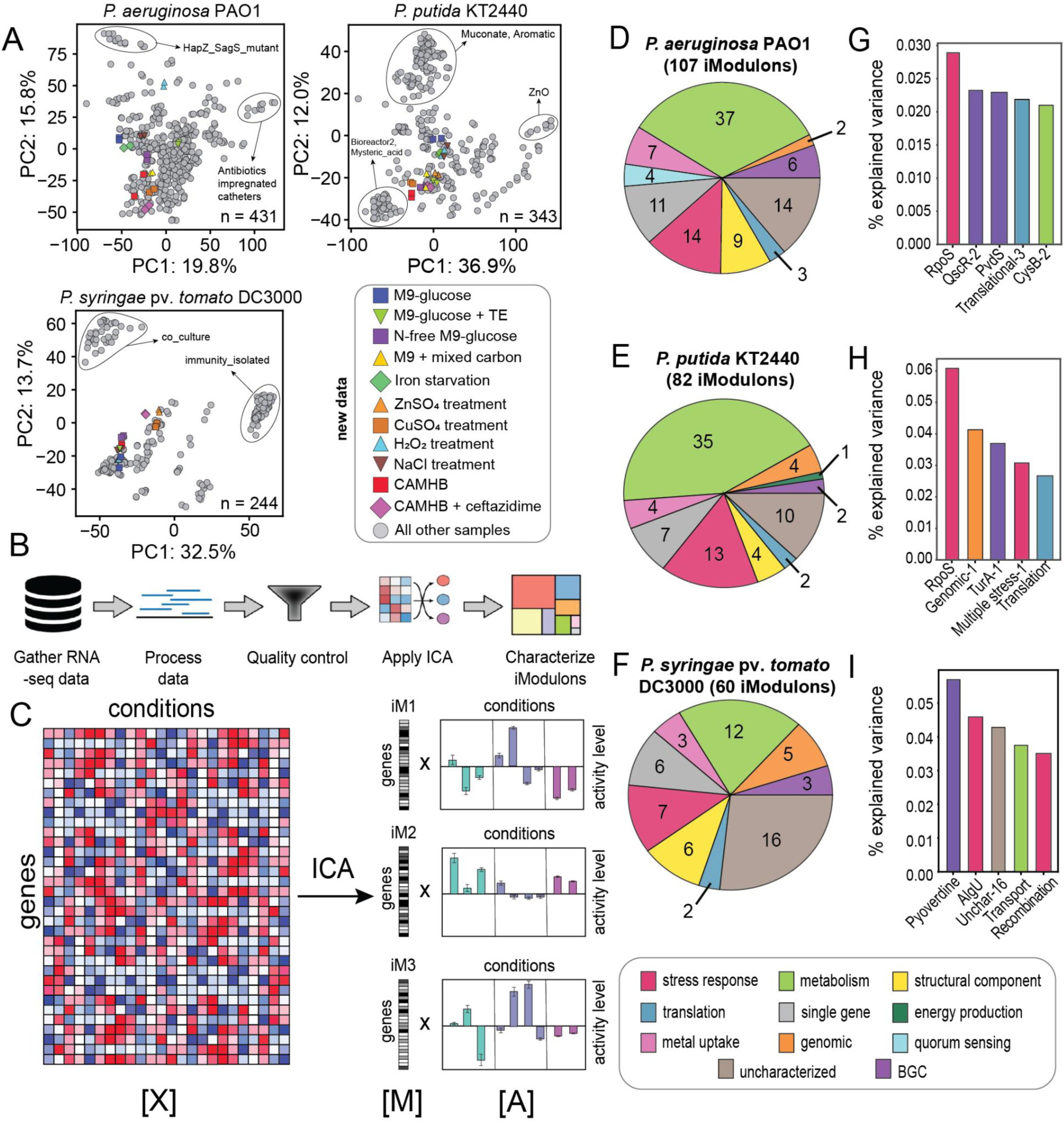
Overview of iModulons of three Pseudomonas strains. (A) Scatter plots of first two principal components of the RNA-seq compendiums for each strain. Colored sample points represent new samples generated for the present study. Samples from select projects have been highlighted and labelled by project name. (n: number of RNA-seq samples in dataset) (B) Data processing pipeline used to run ICA to generate iModulons. This diagram was adapted from Anand et al(20). (C) ICA decomposes the gene expression profiles into an “**M**” matrix containing the weights of each gene in the iModulons identified and an “**A**” matrix containing the activity of each iModulon across all conditions (iM: iModulon). (D-F) Pie charts represent iModulon counts per functional category for each strain. (G-I) Bar graphs show the percentage of variance explained by the top five iModulons with the highest explained variance for each strain. (BGC: biosynthetic gene cluster)

After normalizing log_2_[TPM] data to project-specific reference conditions, we applied ICA to each dataset(17). We extracted 107, 82, and 60 iModulons for *P. aeruginosa*, *P. putida*, and *P. syringae*, respectively (Fig. 1B, Supplementary Table S4-6; Supplementary Note S3). Each iModulon represents a group of genes that are co-regulated within the given condition-space. ICA provides iModulon gene compositions and activities across all conditions (Fig. 1C). iModulons correspond to the effects of transcription factors under conditions within the dataset, thus having both statistical and biological significance.

### iModulons enable genome-scale TRN comparison

After curating and characterizing the iModulons, they were categorized based on regulator enrichment. iModulons that are significantly enriched for a single regulon were categorized as “regulatory”. The remaining iModulons were categorized as “biological” (representing a biological function), “genomic” (representing genomic features), “uncharacterized” (representing unknown cellular functions), or “technical” (containing a single highly weighted gene, likely linked to noise in gene expression). Comparison of regulator enrichment across strains reveals an overall lack in available regulatory annotations in *P. syringae* (Supplementary Fig. S1; Supplementary Note S4).

The iModulon structures of all three strains capture diverse cellular functions, such as metabolism, stress responses, and translation (Fig. 1D-F). “Metabolic” iModulons dominate *P. aeruginosa* and *P. putida*, while “uncharacterized” iModulons dominate *P. syringae* (16). Together with the low count of “regulatory” iModulons identified in *P. syringae*, these findings highlight the lower level of TRN characterization for this strain. The availability of iModulon structures for all three strains provides a unique opportunity to improve characterization of the *P. syringae* TRN through cross-strain iModulon comparison, as detailed in Supplementary Note S5.

Functional categorization of iModulons also reveals strain-specific characteristics. For instance, the “quorum sensing” (QS) category is unique to *P. aeruginosa*, which is known to have multiple QS systems that play key roles in pathogenicity and antibiotic resistance (18). The absence of QS iModulons in the other strains may be due to a lack of conditions differentially activating the associated genes. Alternatively, QS may not be regulated in a coordinated manner in these strains, resulting in ICA being unable to identify any associated signals.

Additionally, the strains differed based on the iModulons capturing the highest percentage of variance in gene expression within each dataset (Fig. 1G-I; Supplementary Note S6). While stress response iModulons capture a high percentage of variance in all three strains, the associated regulators vary. iModulons enriched for RpoS, which governs the general stress response, capture the highest percentage of variance within the *P. aeruginosa* and *P. putida* datasets. However, the primary stress response iModulon in *P. syringae* is enriched for AlgU, which plays a larger role in stress tolerance in the strain compared to RpoS(19). These differences suggest that although stress adaptation is a major driver of transcriptional variation across the strains, the underlying regulatory programs have diverged.

Together, these observations suggest that cross-strain iModulon comparison can reveal conserved and diverged regulatory mechanisms across the strains.

### Cross-strain iModulon mapping identifies conserved and strain-specific regulatory modules

To compare iModulons across the pseudomonads, we first used OrthoFinder to identify orthologous gene groups(21). Of 4,556 orthogroups identified, 2,063 have at least one gene member belonging to an iModulon (Supplementary Table S7; Supplementary Fig. S2; Supplementary Note S7). We mapped iModulons across the strains based on the absolute correlation (Pearson’s *r*) of **M** matrix gene weights of orthologous genes (see Methods) to generate an iModulon network map (Fig. 2A; Supplementary Table S8). We observed that 52% (56/107), 63% (52/82), and 68% (41/60) of iModulons in *P. aeruginosa*, *P. putida*, and *P. syringae*, respectively, are connected to at least one iModulon in another strain based on orthologous gene weights (Pearson’s *r*<0.3). Of these, 14% (15/107), 17% (14/82), and 15% (9/60) of iModulons in *P. aeruginosa*, *P. putida*, and *P. syringae* had a Jaccard similarity of at least 0.5 with at least one iModulon in another strain, representing transcriptional signals involving the expression of a core set of orthologous genes across the strains. Correlated iModulons with high similarity in orthologous gene membership often represent essential cellular functions, such as translation. In contrast, those with low similarity represent evolved functions with notable differences in gene membership due to lineage-specific adaptations, such as stress responses.

**Figure 2.**
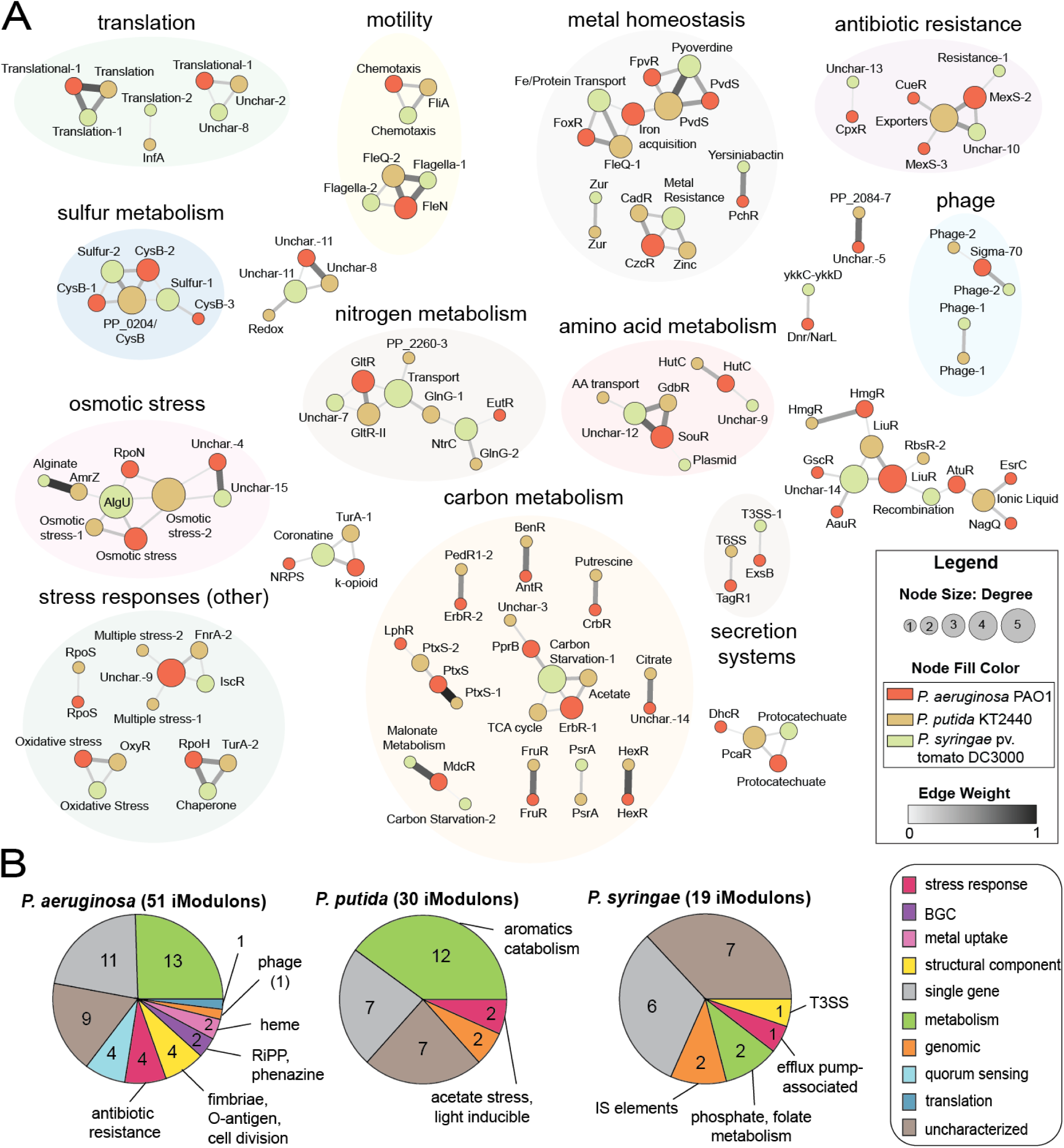
Network map of iModulons. (A) Network map of iModulons categorized based on dominant functional category. (B) Pie charts of unmapped iModulons across the three strains categorized based on iModulon function (IS: insertion sequence; BGC: biosynthetic gene cluster). (PA: P. aeruginosa, PP: P. putida, PS: P. syringae)

Mapping iModulons across strains also highlights strain-specific regulatory mechanisms, represented by iModulons that fail to match to at least one iModulon from another strain (Fig. 2B). While the detection of unique iModulons may be due to condition space differences (Supplementary Note S8), iModulons that represent true unique capabilities of strains were identified. These iModulons often contain genes with no known orthologs within the other strains. For instance, *P. aeruginosa* contains unique iModulons involved in antibiotic resistance and O-antigen biosynthesis which contribute to its virulence and pathogenicity. Additionally, *P. putida* contains unmapped iModulons that correspond to aromatic compound metabolism, representing the persistence of the strain against toxic substances in its natural environment. Finally, several unique iModulons of *P. syringae* can be linked to plant pathogenicity, such as type III secretion system (T3SS) generation. Overall, the iModulon network map serves as a basis for cross-strain TRN comparison.

### iModulons reveal differences in activation of translation

iModulons representing fundamental cellular functions across the strains often share a large core orthologous gene set. Nevertheless, these iModulons may have different activation states in similar conditions. We studied a cluster of translation-associated iModulons with high orthologous gene membership similarity, containing the Translational-1 (*P. aeruginosa)*, Translation (*P. putida)*, and Translation-1 (*P. syringae)* iModulons. Despite sharing 27 orthologous genes encoding ribosomal proteins (Supplementary Fig. S3; Supplementary Note S9), their activities differ within the shared condition space (Fig. 3A). For instance, *P. aeruginosa* and *P. putida* strongly upregulate their translation iModulons in CAMHB media compared to M9 media. Additionally, they downregulate these iModulons during treatment with the cell wall active antibiotic, ceftazidime. These activities reflect the expected pattern of the fear-greed trade-off: faster growth in rich media, and retardation of growth in stressful conditions(22). In contrast, the Translation-1 iModulon of *P. syringae* displayed a lack of increased growth in CAMHB media. Additionally, treatment with ceftazidime strongly upregulated the iModulon. Being a phytopathogen, encountering a stressor is often an indicator of the activation of the plant immune system to *P. syringae*. As a response, the bacteria increases translation of T3SS effectors that suppress the immune system, as seen with the coordinated upregulation of the T3SS-1 iModulon (Fig. 3B). Thus, the upregulation of the Translation-1 iModulon in this stress condition may be a result of divergent adaptations influenced by host-related pressures.

**Figure 3.**
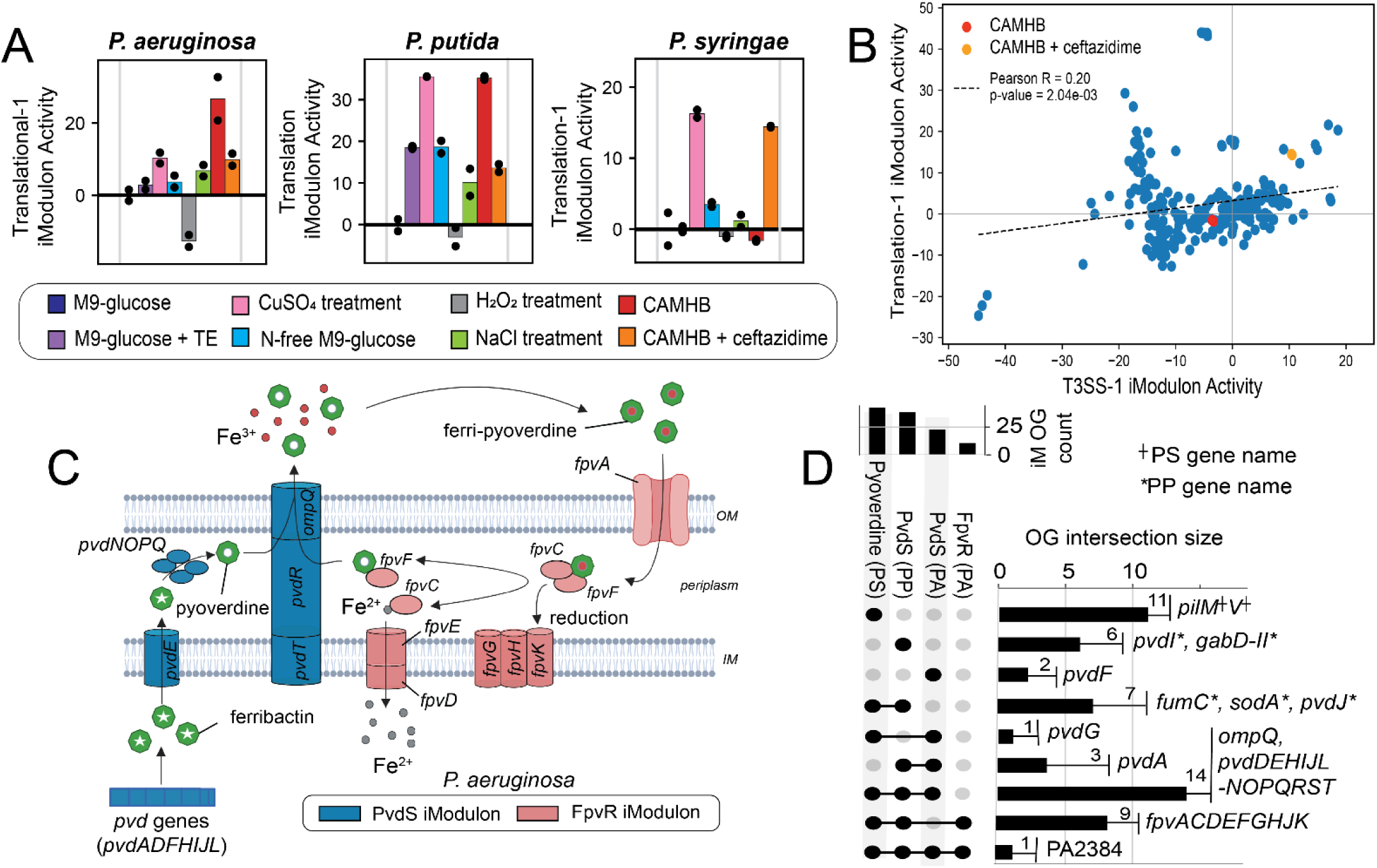
Cross-strain iModulon comparison based on iModulon gene membership and activities. (A) Activity of translation-associated iModulons across common conditions in the new RNA-seq data generated in this study (TE: trace element solution). (B) Scatter plot of the activities of the T3SS-1 and Translation-1 iModulons. (C) General pyoverdine production and uptake pathways in pseudomonads. The pathway is colored based on P. aeruginosa iModulon membership of genes encoding pathway components (OM: outer membrane; IM: inner membrane) (Created using Biorender)(23, 24). (D) Upset plot of orthogroup membership of pyoverdine-associated iModulons. Orthogroup counts include unique genes not assigned to any orthogroup. (PA: P. aeruginosa; PP: P. putida; PS: P. syringae; iM: iModulon; OG: orthogroup)

These results highlight that strong conservation of iModulon gene membership does not necessarily translate to conserved activation states, reflecting divergence in regulatory responses across strains. We also performed a broader analysis of iModulon activities across the strains, details of which can be found in Supplementary Note S10 (Supplementary Fig. S4-5).

### *P. aeruginosa* uniquely regulates pyoverdine biosynthesis independently from pyoverdine uptake

Next, we analyzed iModulons involved in pyoverdine-mediated iron acquisition(25). In iron-limiting conditions, *Pseudomonas* strains produce a fluorescent siderophore called pyoverdine which chelates iron (Fig. 3C)(23). The newly formed ferri-pyoverdine complex is then taken up by the cell and reduced to a usable form. Pyoverdine biosynthesis and uptake are regulated by PvdS and FpvR, respectively. *P. aeruginosa* contains two iModulons individually enriched for the regulons of each regulator, namely the PvdS and FpvR iModulons. In contrast, *P. putida* and *P. syringae* both have a single iModulon representing both pyoverdine biosynthesis and uptake (*P. putida*: PvdS iModulon; *P. syringae*: Pyoverdine iModulon).

To understand the unique split of the PvdS and FpvR regulated genes into two individual iModulons in *P. aeruginosa*, we compared the activities of the iModulons across the dataset (Supplementary Fig. S6). Indirect regulatory mechanisms linked to virulence as well as differences in spatial structures of bacterial growth modes (i.e., biofilm, planktonic, and dispersed cell growth) contribute to divergence in the activities of the otherwise strongly correlated iModulons (Pearson’s *r*=0.71) (Supplementary Note S11). The extraction of individual PvdS and FpvR iModulons in *P. aeruginosa* may be linked to the pathogenicity of the strain and its consequent versatility. Alternatively, this may be a regulatory strategy employed to enable recycling of internalized pyoverdine, thereby reducing the energetic cost of pyoverdine biosynthesis(26).

We also compared gene membership of the pyoverdine biosynthesis and uptake iModulons across strains (Fig. 3D). Most genes in the pyoverdine iModulons of *P. putida* and *P. syringae* have orthologs in either the PvdS or FpvR iModulons of *P. aeruginosa*. However, some genes are unique to the iModulons of the two strains. For instance, seven orthologous genes are shared by the pyoverdine iModulons of *P. putida* and *P. syringae*, but absent in those of *P. aeruginosa*. These include oxidative stress mitigation genes *sodA* and *fumC*, indicating a coupling of oxidative stress and iron limitation responses in the strains. Together, iModulon structures and gene membership patterns underscore both the shared and lineage-specific adaptations in pyoverdine regulation.

### Motility iModulons reveal varying needs across bacterial lifestyles

A cellular function common among pseudomonads is flagella-mediated motility(28). Each of the three strains possesses a primary flagellar assembly iModulon: FleN (*P. aeruginosa*), FleQ-2 (*P. putida*), and Flagella-1 (*P. syringae*) iModulons (Supplementary Fig. S7). *P. syringae* contains a second flagellar assembly iModulon, Flagella-2, which is hypothesized to enable evasion of the plant immune system(14). We found a conserved set of orthologous genes in the primary flagellar assembly iModulons, most of which encode structural components of the flagella (5A-B; Supplementary Fig. S8; Supplementary Note S12).

While the gene neighborhoods harboring these genes are highly conserved, some iModulon genes within these regions are unique to individual strains. The FleN iModulon of *P. aeruginosa* contains a glycosylation island formed by three unique genes, PA1088-PA1090 (Fig. 4A)(29, 30). This BGC is involved in glycan biosynthesis and is linked to virulence and host cell-specific interactions. We also found unique iModulon genes linked to flagellar arrangements. Of the three strains, *P. putida* possesses the highest number of flagella(31). A unique gene within the FleQ-2 iModulon, PP_4331, is predicted to influence flagellation patterns(34) (Fig. 4B). Additionally, while the *fleN* gene involved in flagellar numerical control is found in all three strains, it is not an iModulon member in *P. putida*. This may indicate a lower restriction on transcriptional control of flagellar count, allowing the strain to produce more flagella and move through viscous soil environments it typically inhabits. Overall, these findings suggest that unique iModulon genes may represent key drivers of strain-specific phenotypes despite overall conservation of the genomic regions.

**Figure 4.**
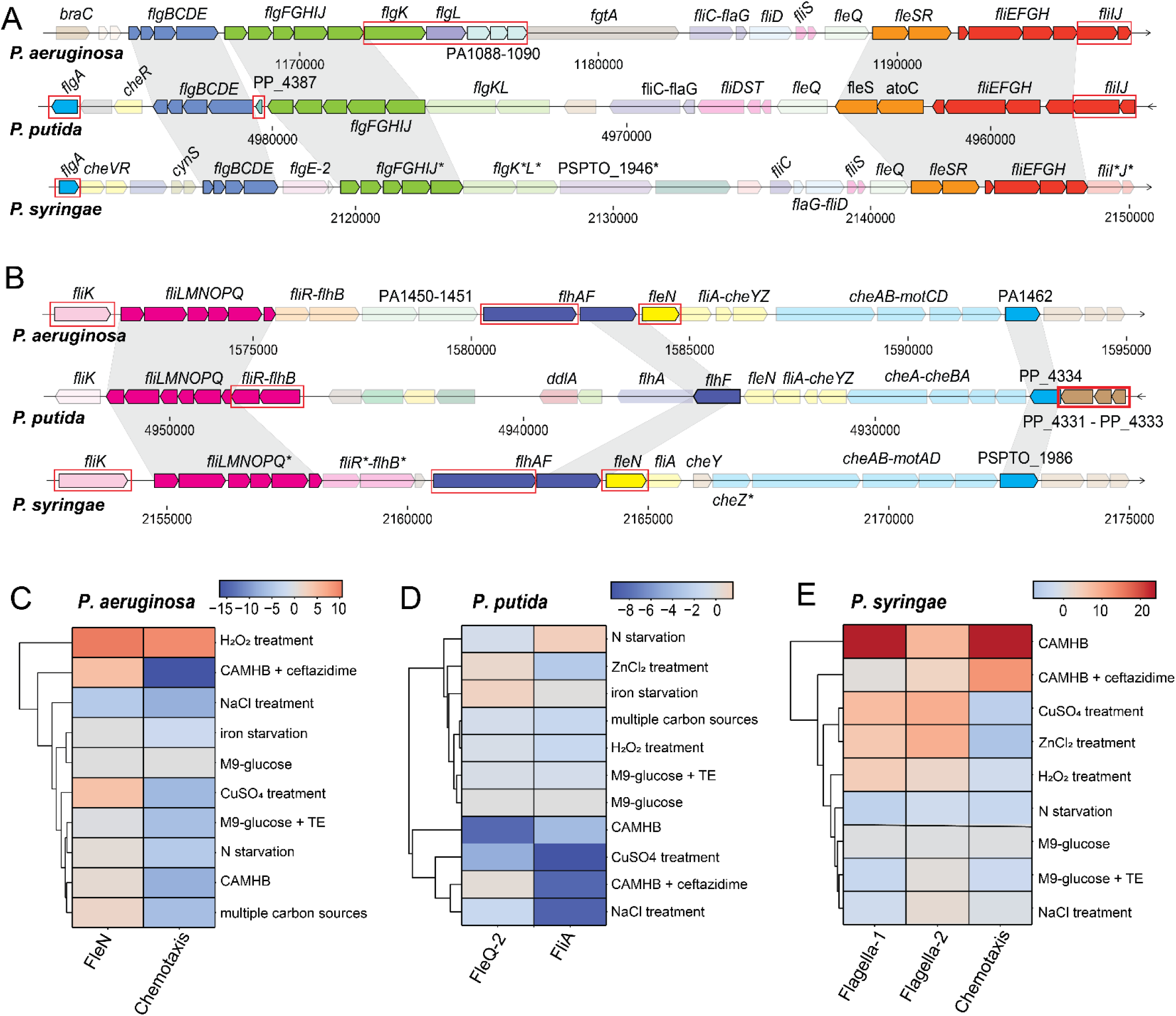
Motility iModulons of Pseudomonas strains. (A,B) Comparison of gene maps for two neighborhoods harboring most gene members of the primary flagella-assembly iModulons of the three strains (P. aeruginosa: FleN iModulon; P. putida: FleQ-2 iModulon; P. syringae: Flagella-1 iModulon). Gene members of these iModulons are in bold colors, and orthologs unique to one or two of these iModulons across all three strains are highlighted in red. Genes found in the Flagella-2 iModulon of P. syringae are marked with an asterisk. (C-E) Heatmaps of activities of motility iModulons in the conditions newly generated in this study. (TE: trace element solution; multiple carbon sources: mixture of succinate, fructose, and toluene)

We also identified differences in motility iModulon activities under common conditions (Fig. 4C-E). *P. syringae* strongly downregulates both flagellar assembly and chemotaxis in the presence of ceftazidime, whereas *P. putida* and *P. aeruginosa* upregulate both functions. Additionally, *P. aeruginosa* and *P. syringae* have stronger responses to oxidative stress and metal ion stress, respectively, in comparison to the other strains. Interestingly, the motility iModulons of *P. putida* have a very narrow positive activity range (Fig. 4D). This may be attributed to the existence of the strain in homogeneous environments where swimming away from stressors is most often not beneficial. Further analysis of iModulon activities may thus reveal similarities or differences in the “flight” component of stress responses across the strains that can be linked to strain-specific biology.

### iModulon activities uncover varying antibiotic stress mitigation mechanisms across strains

Another feature of interest of *Pseudomonas* species is antibiotic resistance (35). Using iModulons, we studied transcriptome-wide responses to ceftazidime treatment (Fig. 5A-C). Multiple carbon and amino acid metabolism-associated iModulons were differentially activated in the ceftazidime treatment condition in *P. aeruginosa* and *P. putida*, although they largely represent different metabolic functions in each strain. Additionally, only *P. syringae* was observed to differentially upregulate all translation-associated iModulons during ceftazidime treatment. As previously suggested, this may be a host interaction-based adaptation.

**Figure 5.**
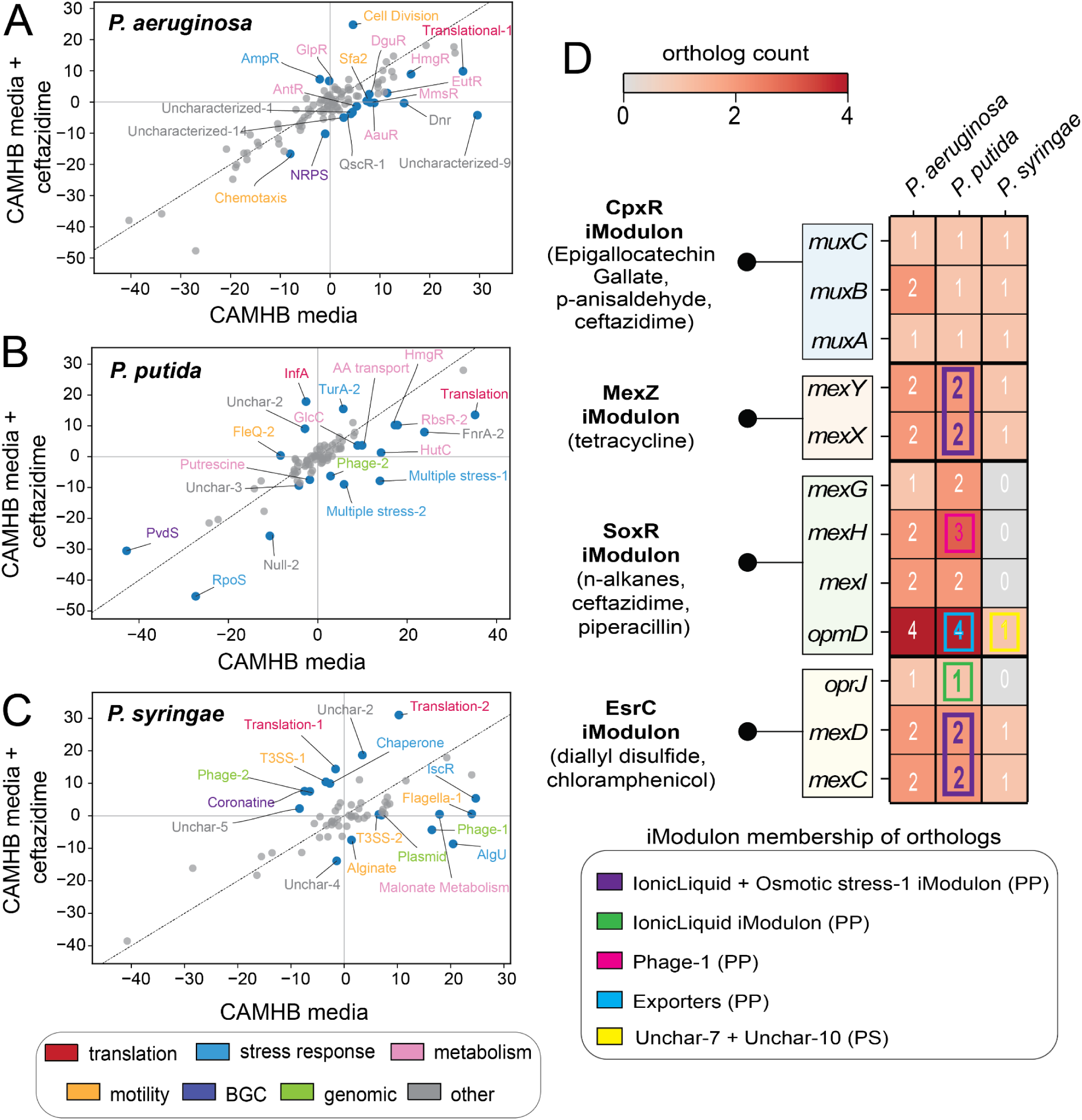
Antibiotic stress responses of Pseudomonas strains. (A-C) Differential iModulon activation plots for the three strains, comparing growth in CAMHB media to growth in CAMHB media treated with ceftazidime. (D) Heatmap depicting the number of orthologous genes across the three strains for select genes within antibiotic responsive iModulons of P. aeruginosa(12). All efflux pump genes within the selected iModulons are depicted. Activating conditions for each selected iModulon are listed in brackets. The iModulon membership of orthologs of selected genes is highlighted in the colored boxed regions, and is indicative of the iModulon membership of at least one of the identified orthologs.

The antibiotic stress responses of *P. putida* and *P. syringae* are global and non-specific. Upon antibiotic treatment, both strains activated non-specific stress response iModulons such as chaperone-associated iModulons. However, iModulon activations in *P. aeruginosa* revealed specialized response components. The only stress response iModulon that was differentially activated in the strain was the AmpR iModulon, containing the beta-lactamase gene *ampC*. Additionally, the most strongly differentially activated iModulon was the Cell Division iModulon, involved in cell wall biosynthesis. Both the AmpR and Cell Division iModulons are unique to *P. aeruginosa*, revealing the unique antibiotic resistance capabilities of the strain.

### Abundance of antibiotic-responsive iModulons contribute to antibiotic resistance in *P. aeruginosa*

To further explore the distinctive antibiotic resistance capabilities of *P. aeruginosa*, we studied previously identified antibiotic-responsive iModulons(12). The AmpR iModulon in *P. aeruginosa* contains four genes (*ampC*, *creD*, PA4111, and PA0466) and is involved in beta-lactam resistance (Supplementary Fig. S9A)(37). While orthologs for *ampC* and PA4111 exist in *P. putida* and *P. syringae*, *creD* and PA0466 lack orthologs in these strains. Additionally, while the expression of the AmpR iModulon genes showcase strong positive correlation with one another across the *P. aeruginosa* dataset, the expression of *ampC* and PA4111 orthologs in *P. putida* and *P. syringae* display low correlation with one another (Supplementary Fig. S9B-D). This indicates a lack of coexpression of these orthologs across their respective datasets, resulting in the absence of an AmpR-associated iModulon.

Bacteria also utilize efflux pumps to resist toxic compounds. *P. aeruginosa* possesses four iModulons containing genes encoding efflux pumps previously found to be antibiotic-responsive: CpxR (*muxABC*), MexZ (*mexXY*), SoxR (*mexGHI*-*opmD*), and EsrC (*mexCD*-*oprJ*) iModulons(12). Efflux pump genes in these iModulons contain at least one ortholog in *P. putida* (Fig. 5D). Several of these orthologs in *P. putida* are iModulon members, and at least one ortholog for the selected genes in the MexZ and EsrC iModulons are members of the IonicLiquid iModulon. This iModulon is strongly responsive to triethylammonium hydrogen sulfate. Future studies may reveal its responsiveness to antibiotics such as those activating the MexZ and EsrC iModulons of *P. aeruginosa*. In contrast to *P. putida*, a few of the selected genes lack orthologs in *P. syringae*. Of the genes with orthologs in the strain, only one was an iModulon member. Since the condition space of *P. syringae* mostly consists of plant infection and co-culture conditions, it is possible that these efflux pumps are not activated in this dataset. With the antibiotic treatment conditions within the *P. aeruginosa* dataset as reference, expansion of the *P. putida* and *P. syringae* datasets may reveal specific antibiotic resistance mechanisms and contribute to a more comprehensive iModulon coverage of efflux pumps. Alternatively, these conditions may elicit responses that are aligned with the previously identified general stress response patterns observed during ceftazidime treatment.

## Discussion

In this study, we present a genome-scale comparison of the TRNs of *P. aeruginosa* PAO1, *P. putida* KT2440, and *P. syringae* pv. *tomato* DC3000. After expanding strain-specific RNA-seq datasets with eight common growth conditions, we applied ICA to each dataset to extract 107, 82, and 60 iModulons for *P. aeruginosa*, *P. putida*, and *P. syringae*, respectively. Mapping iModulons across the strains based on orthologous gene membership reveals shared and unique regulatory modalities. Comparative analysis of iModulon gene membership and activities provides insight into the diversification of TRNs. We find that conserved cellular functions, including translation and pyoverdine biosynthesis/uptake, can undergo strain-specific regulatory adaptations through both differences in iModulon gene membership and condition-specific activation patterns. Additionally, analysis of motility-associated iModulons reveals how transcriptional regulation of conserved processes reflects differences in bacterial lifestyles, including environmental adaptation, virulence, and host-associated behaviors. Finally, genome-level comparisons of antibiotic-responsive iModulons uncover divergent antibiotic stress response strategies across the strains and highlight the enrichment of specialized virulence-associated functions in *P. aeruginosa*.

As an advancement over previous iModulon studies focused primarily on individual strains, this study establishes a scalable framework for comparing TRNs across closely related strains. We first expanded the existing strain-specific transcriptomic datasets by incorporating shared growth conditions. This common experimental framework allows us to compare both TRN structure and regulator activity under common conditions. The improved comparability across the datasets provides new opportunities to investigate strain-specific regulatory strategies. A major finding of this study is that conservation of iModulon structure does not imply conservation of regulatory dynamics. Our analysis suggests that even essential cellular processes like translation can be rewired to support different lifestyles of bacteria. This highlights the importance of comparing both iModulon composition and activity across strains. Furthermore, this framework enables systematic comparisons of global transcriptome compositions across strains exposed to the same environmental pressures. We identified differences in the extent to which motility is employed as a stress response among the three pseudomonads, and demonstrated the utility of this method in enabling a comprehensive genome-scale comparison of stress response mechanisms. These case studies illustrate the breadth of biological insights that can be gained through cross-strain iModulon analysis.

The iModulon structure of a strain provides a modularized representation of the TRN that allows it to adapt to the condition space of the dataset used to compute iModulons. While transcriptional signals like translation are captured by ICA with a minimally diverse condition space, the comprehensiveness of the iModulons obtained depend on the conditions within the dataset. As a result, the comprehensiveness of cross-strain iModulon comparison is improved by increasing the similarity between the condition spaces of the datasets being analyzed. This enables the capturing of transcriptional responses of each strain to a more similar set of activating conditions. In this study, we provide a first step towards building datasets for *P. aeruginosa*, *P. putida*, and *P. syringae* with greater condition space comparability through the addition of eight new overlapping conditions. With the new set of conditions, along with a few others that roughly compare, we show how condition-specific responses can be compared through cross-strain iModulon analysis. Given the scalability of iModulon computation, future studies can further expand on our analysis through sample generation to improve the condition space overlap (Supplementary Note S8).

In conclusion, we have demonstrated the utility of iModulons in comparing the TRNs of strains at the genome scale. By comparing iModulons across *Pseudomonas* strains, we can trace the evolutionary dynamics along their lineages. Altogether, this study provides a roadmap for cross-strain iModulon analysis.

## Methods

### RNA extraction and library preparation

The following strains were used in this study: *P. aeruginosa* PAO1, *P. putida* KT2440, and *P. syringae* pv. *tomato* DC3000. Cells were grown overnight at 30 °C with shaking in 4 mL of the base medium indicated in Supplementary Tables S1-3. The following day, fresh cultures were inoculated in duplicate to an initial OD_600_ of 0.05. For iron starvation (2,2’ dipyridyl treatment) and nitrogen starvation (nitrogen-free M9 media with addition of eighteen amino acids, except for cysteine and methionine) conditions, cultures were grown in medium supplemented as specified in Supplementary Tables S1-3. Cultures were harvested at mid-log phase (OD₆₀₀ ≈ 0.5) and 3 mL of culture was immediately mixed with 6 mL of RNAprotect reagent for the following conditions: growth in M9 glucose media (with and without trace element solution), CAMHB media, mixed carbon source (succinate, fructose, and toluene treatment), iron starvation, and nitrogen starvation. For ceftazidime treatment conditions, ceftazidime (¼ of minimum inhibitory concentration) was added at OD₆₀₀ ≈ 0.2, and cultures were incubated for 1 h prior to collection. For osmotic stress (NaCl treatment), oxidative stress (H_2_O_2_ treatment), and metal ion stress (CuSO_4_ and ZnSO_4_ treatment) conditions, cultures were grown to OD₆₀₀ ≈ 0.5 before addition of the corresponding supplement to the final concentration indicated in Supplementary Table S1-3. This was followed by incubation for an additional 30 min (osmotic and oxidative stress conditions) or 1 h (metal ion stress conditions) before mixing 3 mL of culture with 6 mL of RNAprotect reagent. Cells were pelleted and stored at −80 °C until RNA extraction and RNA-seq library preparation.

Total RNA was isolated and prepared from cell pellets using reagents from Zymo Research and silica-coated magnetic beads (Luna Nanotech). Cell pellets were resuspended in 415 uL RNA Binding Buffer (Zymo Research) containing 8 uL Beta-mercaptoethanol in 1.5 mL tubes. After dislodging pellets into the buffer using an inoculating loop, they were poured into 2 mL tubes containing ∼250 uL 0.1mm diameter zirconia/silica beads (BioSpec Products). Any remaining buffer was transferred by pipetting. Cells were lysed using an Omni International Bead Ruptor 12 instrument (2 cycles, 1 min each at maximum speed of 6 m/s) with a 1 min centrifugation at 10,000 x g between cycles. Lysates were cleared by centrifugation (2 min, 10,000 x g). A total of 200 uL of the cleared lysates were transferred to tubes containing 400 uL 50% ethanol and 20 uL silica-coated beads. Samples were vortexed and incubated at room temperature for 2 min and placed on a magnetic rack until supernatants were clear, after which they were aspirated. Beads were washed once with RNA Wash Buffer (Zymo Research) and resuspended in DNaseI master mix (Zymo Research DNA digestion buffer plus Zymo Research DNaseI). After 15 min incubation at room temperature, each sample was vortexed with 400 uL RNA Prep Buffer (Zymo Research). Samples were returned to the magnetic rack, after which they were washed twice with RNA Wash Buffer and eluted in 15 uL 10 mM Tris, pH 7.5. RNA concentrations and quality were assessed using an Agilent TapeStation and Nanodrop, respectively.

For each sample, 1 ug was used for the standard RiboRid procedure(38). RNA-seq libraries were prepared from 5 uL of the rRNA-subtracted RNA using KAPA RNA HyperPrep Kit (Roche) according to the manufacturer’s protocol with volumes reduced by half. The final PCR amplification step was modified by incorporating SYBR and using a qPCR instrument. Samples were removed from the thermal cycler just before entering the plateau phase of amplification to avoid biases being potentially introduced by reagent depletion. The quality and concentrations of the libraries were checked with an Agilent TapeStation and Qubit instrument, respectively. Pooled libraries were then run on an AVITI instrument (Element Biosciences).

### Data processing and quality control

The iModulonMiner workflow was used to process all candidate RNA-seq samples (Supplementary Note S13)(17). Raw read trimming was performed using Trim Galore (https://www.bioinformatics.babraham.ac.uk/projects/trim_galore/). The reads were then assessed using FastQC (https://www.bioinformatics.babraham.ac.uk/projects/fastqc/) and aligned to the reference genome (Accession numbers: NC_002516.2 (*P. aeruginosa* PAO1); AE015451.2 (*P. putida* KT2440); NC_004578.1, NC_004633.1, NC_004632.1 (*P. syringae* pv. *tomato* DC3000, plasmid pDC3000A, plasmid pDC3000B, respectively)) using Bowtie(39). Read direction was inferred using RSeQC(40), and read counts were generated using featureCounts(41). Finally, quality control metrics were compiled using MultiQC(42). The final expression data are reported in units of log_2_ transcripts per million (log_2_[TPM]).

Samples that failed the following FastQC metrics were discarded: per_base_sequence_quality, per_sequence_quality_scores, per_base_n_content, and adapter_content. Sample with less than 500,000 reads mapped to coding sequences was also discarded. Global correlations (Pearson’s *r*) between the gene expression (log_2_[TPM]) of all samples within each dataset were visualized using hierarchical clustering, and outliers with atypical expression profiles were discarded. Samples without a biological replicate or having low correlation (Pearson’s *r*<0.95) between replicate gene expression (log_2_[TPM]) were dropped. Finally, the log_2_[TPM] data were normalized to the reference condition of each project to remove batch effects. All relevant files can be found on Zenodo (https://zenodo.org/records/18931002).

### Computing iModulons with ICA

We applied ICA to each of the centered log_2_[TPM] datasets by running the Scikit-learn(43) algorithm, FastICA(44), as previously described(10, 17). Using the OptICA method(45), the optimal dimensionality of the *P. aeruginosa*, *P. putida*, and *P. syringae* datasets were determined to be 370, 230, and 200, respectively. The algorithm decomposes a matrix of gene expression profiles (**X**) into an iModulon matrix (**M**) and activity matrix (**A**). The weights of all genes within each component extracted by ICA is provided in the **M** matrix, whereas the activity of each component across all samples in the dataset is provided in the **A** matrix.

The gene weights in the **M** matrix specify the association of each gene with a particular iModulon, with higher absolute weights indicating greater association. The gene membership of an iModulon is defined by applying a threshold to segregate genes based on their weights in an iModulon. These thresholds were calculated for each iModulon using the D’Augustino’s *K*_2_ test for normality, as previously described(10).

### iModulon annotation and curation

The TRNs of all three strains were annotated using RegPrecise(46), Pseudomonas Genome Database(47), and literature. Fisher’s exact test with Benjamini-Hochberg false discovery rate (FDR) correction was used to compute regulator enrichment against known regulons (FDR = 10^−5^). iModulons were then named based on enriched regulators or functional characteristics, with further details in Supplementary Table S4-6.

### Cross-strain iModulon mapping

We ran Orthofinder(21) on default settings to identify orthologous genes. The many-to-many Orthofinder results were used for further steps as suggested on the Orthofinder GitHub page. If more than one gene from a single strain was mapped to an orthogroup, the highest absolute gene weight was selected as the representative gene weight for the orthogroup within an iModulon. The **M** matrix for each strain was reconstructed to provide the weight of an orthogroup in an iModulon, using which iModulons were mapped across the strains using the PyModulon package (cutoff=0.3)(17). After identifying iModulons with Pearson’s *r* correlation of **M** matrix orthogroup weights higher than the selected threshold, we compared iModulons based on orthologous gene membership similarity using the Jaccard similarity metric.

### Differential iModulon activation

Differential iModulon activities between conditions were calculated as previously described(10). The statistical significance of the differences in the activities was determined by comparing the absolute difference in mean iModulon activities between the conditions and the log-normal distribution of the iModulon. iModulons with a threshold > 5 and FDR below the selected value were classified as differentially activated.

## Data Availability

All code and data (aside from raw RNA-seq files) used to generate the results in this paper can be found on Zenodo (https://zenodo.org/records/18931002). New RNA-seq data reported in this study have been deposited in the Gene Expression Omnibus under the accession codes GSE322682, GSE322955, and GSE322719. The general iModulon analysis pipeline can be found at https://zenodo.org/records/18238197.

## Acknowledgements

We acknowledge the National Energy Research Scientific Computing Center for support with high-performance computing.

Author contributions: H.B. and B.O.P designed research. Y.H. collected the experimental data. R.S. and J. Sung performed RNA isolation and library construction. H.B. processed the data, performed the analysis, and wrote the manuscript. B.O.P. provided mentorship and guidance throughout. All participated in reviewing the manuscript.

## Funding

This work was supported by the Novo Nordisk Foundation (NNF) Center for Biosustainability (CfB) at the Technical University of Denmark (NNF20CC0035580) and the Y.C. Fung Endowed Chair in Bioengineering at the University of California, San Diego.

